# Context-dependent choice and evaluation in real-world consumer behavior

**DOI:** 10.1101/2022.04.14.488290

**Authors:** A. Ross Otto, Sean Devine, Eric Schulz, Aaron M. Bornstein, Kenway Louie

## Abstract

A body of work spanning neuroscience, economics, and psychology indicates that decisionmaking is context-dependent, which means that the value of an option depends not only on the option in question, but also on the other options in the choice set—or the ‘context’. While context effects have been observed primarily in small-scale laboratory studies with tightly constrained, artificially constructed choice sets, it remains to be determined whether these context effects take hold in real-world choice problems, where choice sets are large and decisions driven by rich histories of direct experience. Here, we investigate whether valuations are context-dependent in real-world choice by analyzing a large restaurant rating dataset (Yelp.com) as well as two independent replication datasets which provide complementary operationalizations of restaurant choice. We find that users make fewer ratings-maximizing choices in choice sets with higher-rated options—a hallmark of context-dependent choice— and that post-choice restaurant ratings also varied systematically with the ratings of unchosen restaurants. Furthermore, in a follow-up laboratory experiment using hypothetical choice sets matched to the real-world data, we find further support for the idea that subjective valuations of restaurants are scaled in accordance with the choice context, providing corroborating evidence for a general mechanistic-level account of these effects. Taken together, our results provide a potent demonstration of context-dependent choice in real-world choice settings, manifesting both in decisions and subjective valuation of options.

## Introduction

A central theme in the study of decision-making is the limited-capacity nature of human information processing (1). A classic example of choice behavior shaped jointly by presumed processing limitations and the demands of the decision environment is the context-dependent nature of preferences: the relative value of an option depends not only on the option in question but the other options in the choice set, or context (2–5). This sort of context-dependent valuation explains a number of interesting patterns of choice both in humans (6–10) and animals (11, 12).

Context-dependent—rather than stable—valuation famously challenges classical theories of rational choice (13) which posit that value is a static quantity independent of the context, and accordingly, preference between two options should be independent of contextual factors such as the other available options. At the same time, relative (versus absolute) valuation could be understood, computationally, as a rescaling of reward values, relative to the other values present in the context, allowing an individual’s representation of value to adjust to the current value context (6). Indeed, recent theoretical treatments have argued that contextual—versus absolute— value representations permit a decision-maker to discriminate between two low-value options in one context, and at the same time, two high-valued options in another context (14, 15).

Context effects have been observed in both real-world choices—with respect to the temporal history of information viewed, or past choices faced by decision-makers (16, 17)—and in tightly constrained, small-sample demonstrations of “independence of irrelevant alternatives” (IIA) violations (18, 19). While a body of work has examined context effects arising from choice set composition—probing how the distribution of option values faced by the decision-maker systematically alter choices and subjective valuations of options—limited work has investigated whether these sorts of context effects hold in real-world settings (20). Unlike tightly-controlled laboratory settings, the choice sets facing real-world decision-makers are arbitrarily large, and composed of options which could be valued along multiple dimensions, each learned via rich histories of direct experience. Understanding how these real-world context effects unfold is of clear practical importance, as well as for understanding the basic mechanisms of context effects more generally. For example, it is possible that these sorts of context effects do not scale to more naturalistic settings, where individuals may resort to heuristics (e.g. ‘taking the best’). Supporting this view, more naturalistic presentation of options has been shown to reduce apparent suboptimalities in choice (21).

More generally, our understanding of context effects in choice might be inherently limited by the artificial, but well-constrained nature of tasks employed in these laboratory choice studies (22–25). Accordingly, we reasoned that we could leverage large-scale datasets that reflect real-world choices in diverse settings in order to glean novel insights about how context effects play out ‘in the wild’ (26, 27), thus directly evaluating whether these effects are meaningful and general (28). Here we elucidate whether context effects could be observed in actual choices with real-world consequences, for both the decision-maker and the restaurant, at a scale and breadth sufficient to permit detection of context effects while averaging over potential option- and subject-specific effects.

We demonstrate the generality of context-dependent valuation—both in choices and selfreported valuation—by analyzing a massive real-world restaurant rating dataset. Here we examine user behavior on Yelp, a web-based application which maintains crowd-sourced reviews of local businesses which is particularly popular in North America. Our analysis specifically focused on restaurants because, in most cases, people visit one restaurant per meal— ensuring that options effectively compete with each other— and the categories of restaurants are relatively well-defined in our dataset. With the presumption that each Yelp user’s rating evidences both a decision made to visit the rated restaurant over some competing set of options (29) and the user’s subjective valuation of their consummatory experience (30), this rating data permits large-scale examination of contextual effects on both inferred choices and valuations. In our analysis, we view each restaurant decision as one made from a “choice set”—defined as similar, geographically proximate restaurants—with the goal of choosing the restaurant with the highest utility (2, 31–33).

Our large-scale real-world analysis of ratings addresses two empirical questions. First, what is the impact of the local value context upon choice efficiency—namely, do Yelp users make fewer ratings-maximizing choices in sets with higher overall ratings? Based on laboratorybased results indicating that choice contexts increasing the value of ‘distractor’ items (i.e., options lower in value than the higher-rated options in the set) decreased individuals’ propensity to choose the highest-valued item (22, 23)we hypothesized that we would see fewer choices of the highest-rated option in higher-valued contexts (see (22, 24, 34) for discussion of other possibly coexisting effects). One mechanism underpinning such a choice context effect would be if options were selected on the basis of expected values, which are themselves context dependent. To shed light on these intermediate computations we performed a second, laboratorybased study, testing the hypothesis that if decision-makers’ subjective valuations are context dependent, then their ratings of the expected quality of a given restaurant should also exhibit context-dependence in accordance with the value distribution of the choice set, even in the absence of explicit choice. Taken together, we find evidence for context-dependent decisionmaking and valuation in real-world behavior (manifesting both in real-world restaurant choices and ratings) and furthermore, laboratory-based evidence suggesting that these context effects in choice and subjective valuation are closely related.

## Results

We examined signatures of context-dependent choice and subjective valuations using a dataset from Yelp, a website and smartphone app popular in North America, on which users review local businesses such as restaurants and retail stores. Each rating is made on a scale of one to five stars, with higher star ratings indicating a better rating. For every Yelp user considered, we analyzed each user-provided restaurant rating—which, following previous work (29, 31), we took as a proxy for a user’s choice to visit that restaurant at the time of the review— with respect to the competing restaurant options in that neighborhood (i.e., the choice set).

This choice set was defined as all restaurants belonging to the chosen restaurant’s category within the same neighborhood (e.g. “Pizza” in a particular neighborhood in Pittsburgh). These geographic neighborhoods were defined by a density-based spatial clustering algorithm (35, 36), taking all user reviews as input (Figure 1A; see Methods). Thus, the 8 cities considered contained multiple geographically-defined choice sets for each restaurant category which varied considerably in their rating distributions (Figures 1B and C). Taking this approach, we can analyze each user’s choices (N=53,118 choices) in the context of similar restaurants that were geographically proximate (37, 38). Critically, this analysis considered each option’s average rating at the time of the user’s presumed choice—rounded to the nearest half-star, as is presented to users on Yelp— in order to capture the decision-maker’s information state at the time of restaurant selection.

**Figure 1.**
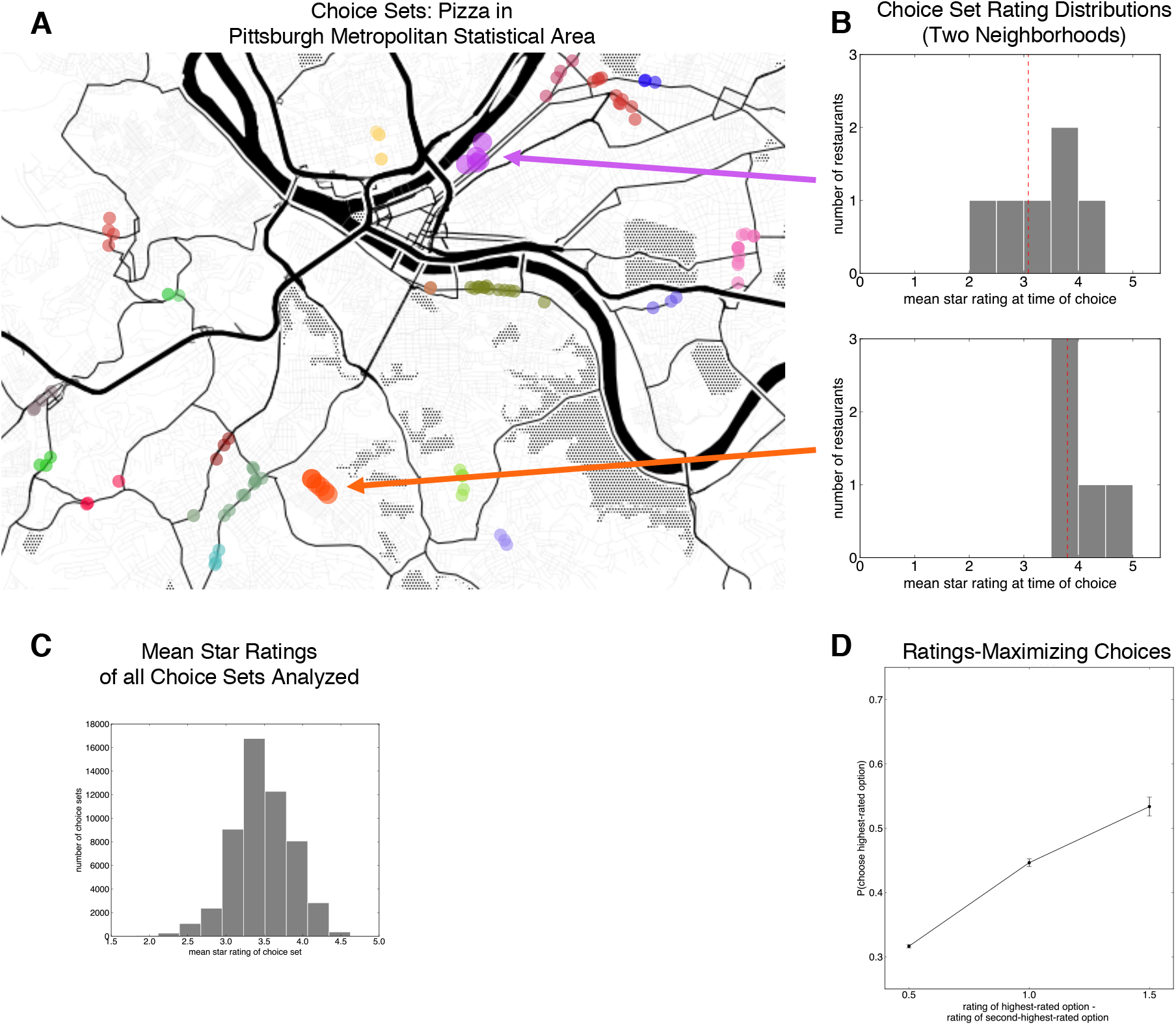
(A) Example of spatially-defined choice sets for Pizza restaurants in Pittsburgh, as computed by the density-based clustering algorithm. Each choice set is denoted by a unique color. (B) Distributions of option ratings for two example choice sets in the Yelp dataset, which correspond to the orange and purple clusters on the map. Notably, the mean star rating (red dashed line) is different across these two choices sets. (C) Distribution of the mean star rating of the choice set, across all choice sets analyzed (D) Proportion of ratings-maximizing (or ‘target’) choices as a function of the difference in ratings between the highest-rated and second-highest-rated option, demonstrating that users were sensitive to the sensitive to option’ ratings.

We first sought to establish that users’ choices were at all sensitive to the options’ ratings. Consistent with the idea that our operationalization of choice and definition of choice sets capture a value-comparison process, the likelihood that a user chooses the highest-rated option (e.g. a 4.5-star option) increased as a function of the difference between the highest-rated option and the second-highest-rated option (e.g., a 4-star option) in the choice set (Figure 1D; logistic regression *β*= 0.5182, 95% CI: 0.4145-0.6149, *p*<0.0001; Supplementary File 1B).

We next examined whether the value context—i.e. the star rating composition of the choice set—affected individuals’ propensity to choose the highest-rated restaurant. We found that as the mean star rating of the choice set increased, choices of the highest-rated option (i.e., target) decreased markedly (Figure 2A, logistic regression *β*=-0.0846, 95% CI: −0.1425, −0.0181, *p=0.002;* Table 1). In other words, the likelihood that a user chose the best-rated “target” depended strongly upon the rating composition of the choice set, with the target choice becoming less likely as the quality of the other options in the set increased, demonstrating a pronounced context effect (5, 23). Intuitively, higher-rated targets were more likely to be chosen (29), and in choice sets with more options, the choice of the highest-rated option was less likely (log number of options *β*=-1.0066, 95% CI: −1.0776, −0.9454, *p*<0.0001), reflecting the well-documented set size effect (34, 39). Importantly, our analysis controlled for the number of reviews (40) and the price of the highest-rated option (31) by taking them as covariates in the regression. Furthermore, this context effect—that, the negative effect of mean choice set star rating upon ratings-maximizing choice—was statistically significant even when controlling for the variance of choice set star ratings (which itself exerted a significant positive effect upon ratings-maximizing choice; *β=* 0.5727, 95% CI: 0.4926, 0.6557, *p*<0.0001; Table 1).

**Figure 2.**
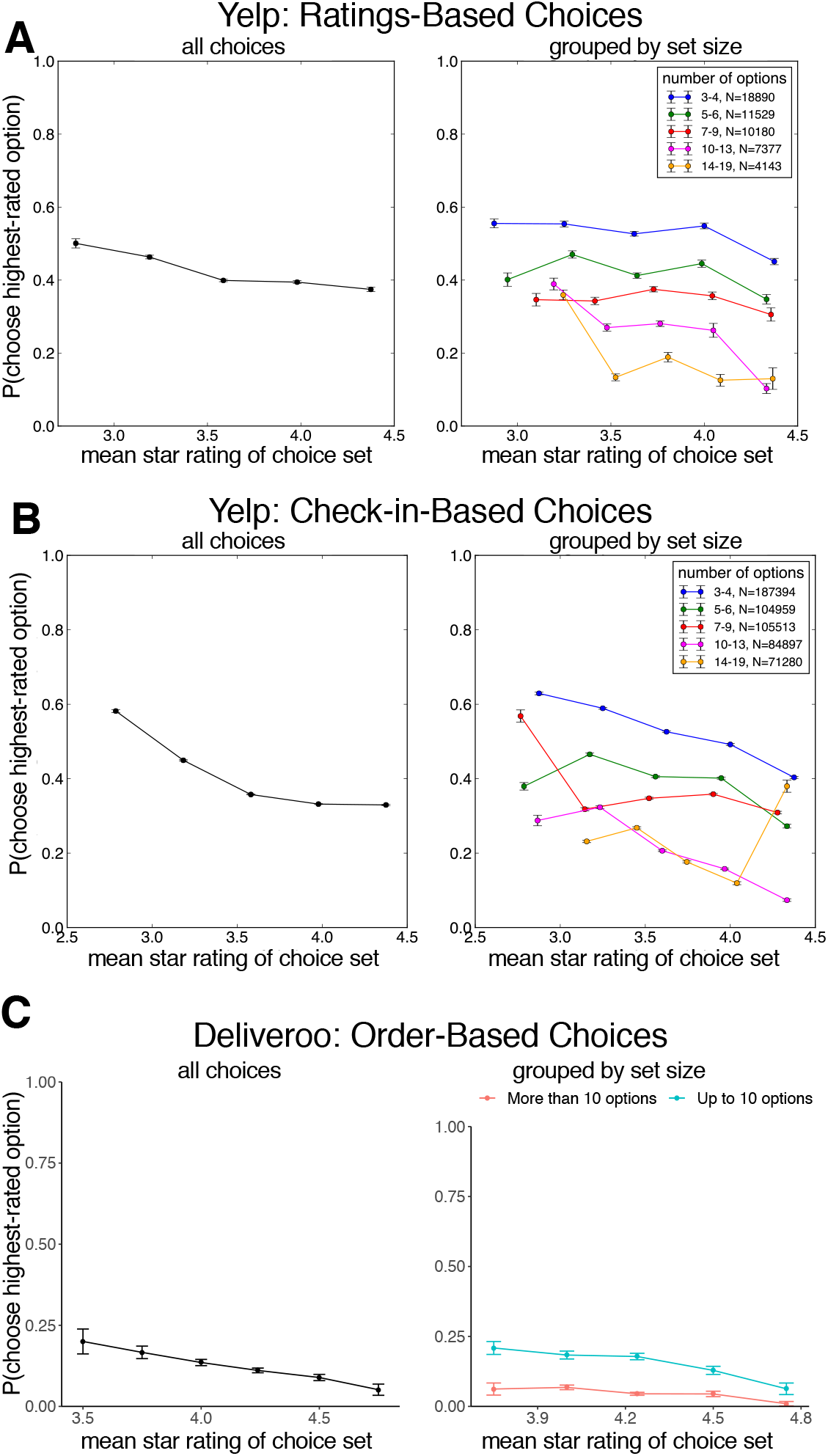
(A) Proportion of choices to the highest-rated option, inferred from Yelp ratings, as a function of the mean star rating of the choice set, for all choice sets (left panel) and broken down by choice set size (right panel). (B) Proportion of choices to the highest-rated option, inferred from Yelp Check-ins, as a function of the mean star rating of the choice set, for all choice sets (left panel) and broken down by choice set size (right panel). (C) Proportion of choices to the highest-rated option, determined from food orders on Deliveroo, as a function of the mean star rating of the choice set, for all choice sets (left panel) and broken down by choice set size (right panel). Across all three datasets, we observed a marked context effect such that participants became less likely to make ratings-maximizing choices as the mean rating of the choice set increased.

**Table 1.**
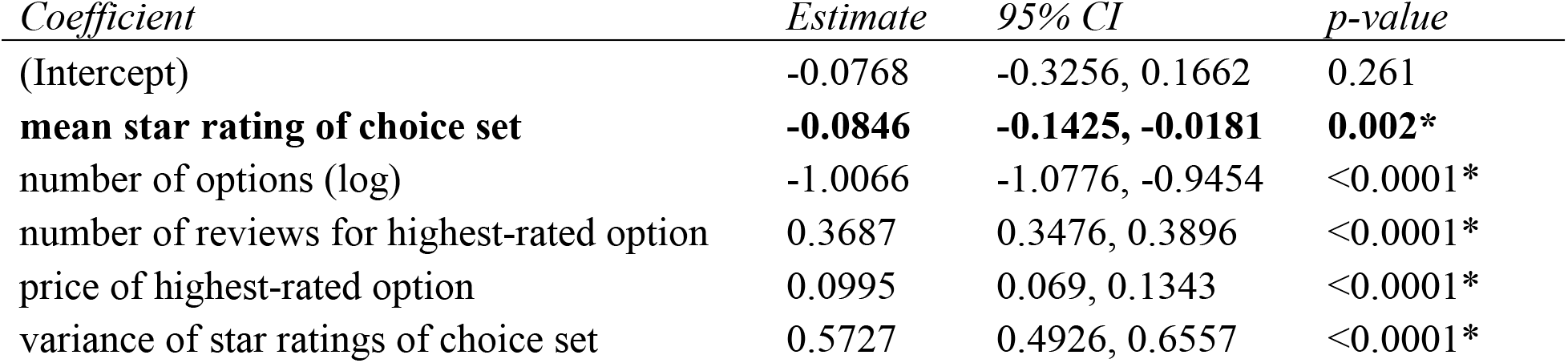
Coefficient estimates for mixed-effects logistic regression predicting ratings-maximizing choices as a function of mean star rating of choice set (i.e., the context effect) and log-transformed choice set size, controlling for the price of the highest-rating option, and the log-transformed number of reviews of the highest-rated option in the Yelp choice dataset.

| *Coefficient* | *Estimate* | *95% CI* | *p-value* |
| --- | --- | --- | --- |
| (Intercept) | -0.0768 | -0.3256, 0.1662 | 0.261 |
| **mean star rating of choice set** | **-0.0846** | **-0.1425, -0.0181** | **0.002\*** |
| number of options (log) | -1.0066 | -1.0776, -0.9454 | <0.0001* |
| number of reviews for highest-rated option | 0.3687 | 0.3476, 0.3896 | <0.0001* |
| price of highest-rated option | 0.0995 | 0.069, 0.1343 | <0.0001* |
| variance of star ratings of choice set | 0.5727 | 0.4926, 0.6557 | <0.0001* |

As we observed a modest correlation between the difference between the top two highest-rated options and the mean star rating of the choice set (Spearman *ρ*=-0.1717, *p*<0.0001), we sought to rule out the possibility that the apparent suboptimality in making ratings-maximizing choices in sets with higher average option ratings (Figure 2A) simply arose from highly-rated second-best options. To do this, we repeated the regression, taking the difference between the ratings of the highest and second-highest options—demonstrated above to predict choice accuracy (Figure 1D)—as a covariate, finding that the effect of mean star rating of choice set (i.e., context) remained statistically significant *(β=* −0.1175, 95% CI: −0.186, −0.0611, *p*<0.001). Further, to address the possibility that the apparent difficulty in making value-maximizing choices in choice sets with higher average option ratings (e.g. Figure 2B) stems from confusion in discriminating the best option, we repeated the analysis of choice as a function of mean choice set rating, only considering choice sets where the second-best option is at least 1 star away from the highest-rated option (24% of choice sets; for example, a set with a highest-rated option of 4.5 stars where the mean rating of the next-best option is 3.5 or less). We found that the contextual effect of mean choice set rating upon ratings-maximizing “target” option selection rates was even stronger in this restricted dataset (*β*= −0.9001, *SE*=0.065, *p*<0.0001) suggesting against the possibility that these context effects arise from ‘crowding’ of options at the higher end of the rating scale. Finally, to address the possibility that high-rated targets exert undue influence on our computed mean choice set rating, we tested if target choices could be predicted as a function of the mean choice set rating without the highest-rated (target) option, finding that the same context effects take hold (*β*= −0.284, *SE*=0.0346, *p*<0.0001).

Importantly, this analysis takes a user’s rating as a proxy for their choice to visit a restaurant, but we sought to demonstrate that these choice context effects take hold in two independent replication datasets which operationalize choice differently. First, we examined a larger dataset of ‘Check-ins’ on Yelp, which are time-stamped, geolocated records of a user having visited a particular restaurant, using the same choice set definition as the previous analysis (N= 565,661 choices; see Methods). Examining users’ propensity to ‘Check into’ the highest-rated option as a function of the mean star rating of the choice set (Figure 2B), we found a similar and pronounced context effect: users were significantly less likely to make choices to the highest-rated ‘target’ option as the quality of the other options in the set increased (*β*= 0.2895, 95% CI:0.223, 0.3672, *p* < 0.000; Supplementary File 1C). We also found that, unlike the ratings-based choice examined above, the variance of the star ratings distribution exerted no discernable effect upon ratings-maximizing choices (*β*=-0.0169, 95% CI: −0.0771, 0.0378, *p*=0.29), suggesting that users’ tendency to make poor choices in choice sets with high mean star ratings purely from a preponderance of higher-valued options in these sets.

Following previous laboratory-based work (23, 24), we performed an additional analysis of choices in the Yelp datasets (inferred from Ratings and Check-ins), examining the ratio of choices made between the two top-rated options as a function of the quality of the “distractor” items inferior to the two highest-rated options. This analysis provides a more direct examination of possible violations of independence of irrelevant alternatives (IIA), according to which the choice ratio between two options should not depend on the quality of other “irrelevant” options (41). Importantly, this analysis only considered the subset of choices made to the highest- and second-highest-rated options in each choice set (75% and 72% of choices in the ratings-based and Check-ins-based choice datasets, respectively), probing whether the relative ratio of choices made to the best option varied as a function of the average ratings of the distractor items rated worse than the best two options. Conceptually replicating Louie et al. (23) and echoing the results of our prior analysis, we observed that choice sets with more valuable inferior options led to significant decreases in the relative ratio of ratings-maximizing choices in both ratings-based choices (*β*= −0.482, 95% CI: −0.5566, −0.4, *p*<0.0001; Figure 2 – Figure supplement 1; Supplementary File 1D) and in Check-in based choices *(β=* −0.497, 95% CI: −0.5131, −0.4802, *p*<0.0001; Figure 2 – Figure supplement 2; Supplementary File 1E). In other words, we observed that users’ ability to choose between high-quality restaurants was systematically diminished by the quality of putatively irrelevant, lower-quality options, a hallmark of context-dependent decision-making.

Further, we examined whether context effects could be observed in actual purchase decisions from the online food delivery service Deliveroo (N=1,613,968 orders; see Methods), a dataset recently used to understand exploratory behavior in consumer choice (31). As in the two other Yelp datasets, we again found a marked context effect (Figure 2C): users became less likely to order from the highest-rated restaurant in a choice set as the average rating of the choice set increased (*β*=-0.01777, 95% CI:-0.0256, −0.0099, *p*<0.001; Supplementary File 1F) but again, variance of star ratings exhibited no significant effect upon choice *(β=* −0.0015, 95% CI: – 0.00920, 0.00627, *p*=0.711). Taken together, these two independent replication datasets support the generality of these context effects of across different settings and operationalizations of choice.

Interestingly, the Yelp ratings examined here—which reflect a user’s consumption experience with a particular restaurant—also afford the opportunity to probe how choice context modulates a user’s subjective valuation of an option following a presumed choice. Intuitively, a user’s restaurant ratings correlated with the average star ratings left by other users before the current rating was made (*r*=0.175, *p*<0.0001), indicating that higher-rated restaurants were more likely to receive higher star ratings after being chosen. Taking this dependency into consideration, we focused on predicting the deviation between the user’s rating and the average restaurant rating at the presumed time of choice (Figure 3A). In other words, we asked whether context had a reliable influence on how a user’s ratings differed from the average ratings, which were left prior that user’s rating. We observed that higher average option ratings of the choice set—i.e., higher-valued contexts—were associated with larger rating deviations (Figure 3B; *β*= 0.0576, 95% CI: 0.037, 0.0753, *p*<0.0001). Importantly, our regression model controlled for a number of variables including the number of previous reviews, the price of the chosen option, and average previous rating of the chosen restaurant (Table 2). At first blush, the directionality of this context effect appears at odds with the context effects observed on choice (Figures 2A-C) where more valuable (or, higher-rated) contexts reduce overall choice efficiency. Following influential accounts of context effects on choice (3, 19, 22, 42), which propose that the subjective value of an option is scaled relative to the values of set of options faced, one might expect a decrease in rating deviation with more valuable (higher-rated) contexts.

**Figure 3.**
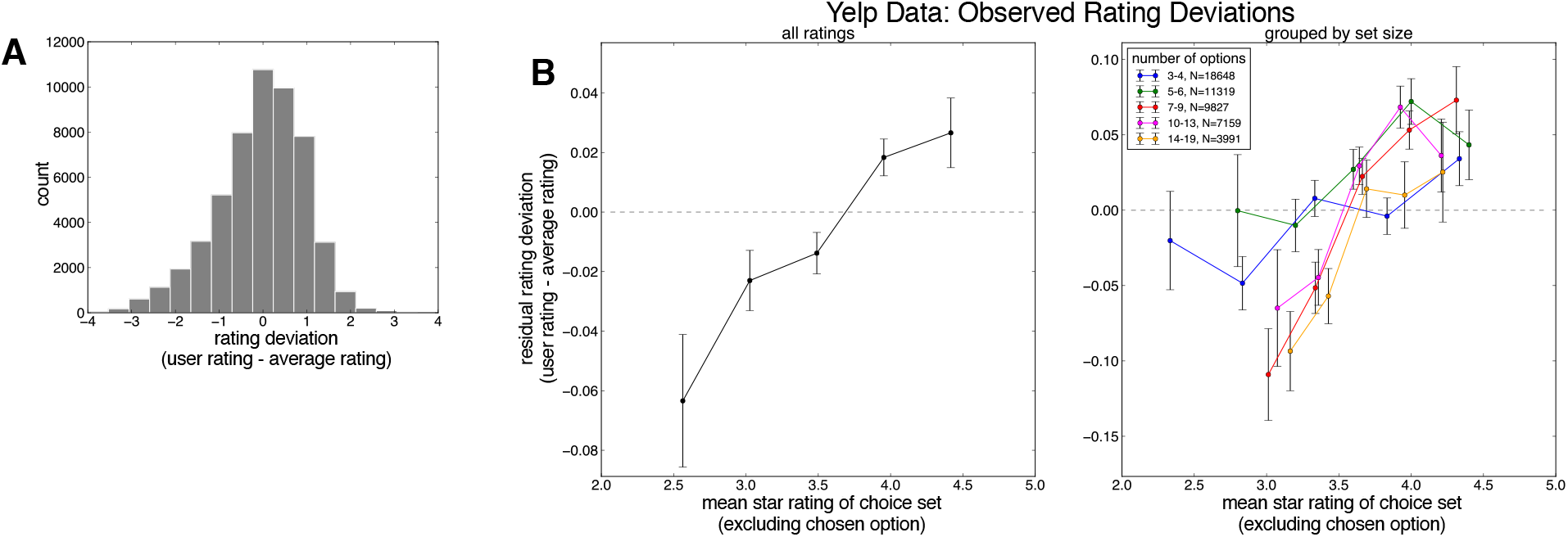
(A) Distribution of deviations between the user’s rating and the average rating of the chosen option at the time of choice for all ratings analyzed. (B) Deviation between the user’s rating and the average rating of the chosen option as a function of the mean star rating of the choice set for all choice sets (left panel) and as a function of choice set size (right panel), after controlling for a number of nuisance variables (see main text). We observed a marked context effect such that higher-rated choice sets resulted in a more positive rating deviation.

**Table 2.** Coefficient estimates for mixed-effects regression predicting the difference between user rating and the mean star rating of the chosen option at the time of choice (rating deviation in Figure 3) in the Yelp dataset, as a function of mean star rating of choice set excluding the chosen option (i.e., the context effect), controlling for the rating of the chosen option at the time of choice log-transformed choice set size, the price of the chosen option, and the log-transformed number of reviews of the chosen option.

| *Coefficient* | *Estimate* | *95% CI* | *p-value* |
| --- | --- | --- | --- |
| (Intercept) | 2.2507 | 2.1255, 2.3526 | <0.0001* |
| mean star rating of chosen option | -0.4338 | -0.448, -0.4197 | <0.0001* |
| length of user review | -0.1759 | -0.1903, -0.1637 | <0.0001* |
| number of reviews for chosen option | 0.027 | 0.0208, 0.0333 | <0.0001* |
| price of chosen option | 0.0843 | 0.0711, 0.099 | <0.0001* |
| **mean star rating of choice set** | **0.0576** | **0.037, 0.0753** | **<0.0001\*** |
| number of options (log) | 0.003 | -0.0136, 0.0211 | 0.368 |

However, it is possible to view ratings not as a pure reflection of the user’s consumption utility of the restaurant—i.e., how good or bad the restaurant was, in absolute terms—but instead as the outcome of a comparison with the user’s contextually bound expectations about the restaurant’s quality (43, 44)before making a rating (and presumably before visiting the restaurant). More formally, we propose that the subjective valuation of an option, reflected in a user’s rating of that option, is computed as the difference between the experienced utility of the restaurant and the contextually-computed expectation of that option (akin to a “prediction error” (45, 46)). On this view, choice contexts with higher average ratings drive should reduce the user’s expectation of an option, in proportion to the average rating of the choice set. Assuming that the user’s experienced utility for an option is a symmetric random variable centered around that option’s ground-truth utility (here, its average rating before the user’s rating is made), an option rated in a higher-valued context should receive higher ratings relative to the options’ ground-truth value. Indeed, this exact pattern manifests in the rating deviations (Figure 3B): more valuable choice sets increase the deviation between a user’s subjective valuation of an option and the average rating of an option.

However, a limitation inherent to this real-world rating data is that only the user’s final rating is observable, but not the user’s contextually computed expectation of a restaurant’s quality, which is a key component in this presumed comparison process. To demonstrate more conclusively that the context effects observed in real-world ratings stem from contextually-bound expectations, we conducted a laboratory experiment in which participants (N=62) viewed choice sets drawn from the real-world choice sets designed to emulate aspects of the choice and rating experience faced by Yelp users (Figure 4A; see Methods). In half of the 270 trials, participants made hypothetical restaurant choices in a given choice set (e.g., pizza in a particular neighborhood), and in the other half of trials, participants provided their expectation of the quality of a randomly selected restaurant on the basis of its restaurant’s rating and the choice set.

**Figure 4.**
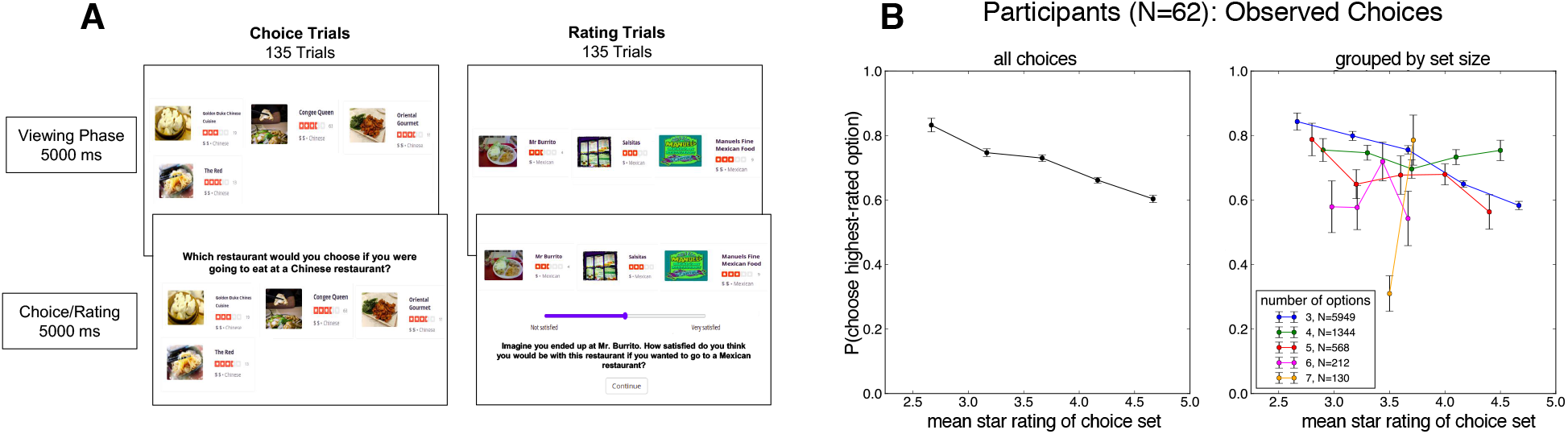
(A) Task screenshots for laboratory experiment. On each trial, participants views a choice set consisting of 3-7 options from a single restaurant category, and either chooses a restaurant (Choice trials; left panel) or indicates their expected satisfaction of a randomly selected restaurant (Rating trials; right panel). (B) Proportion of choices of the highest-rated option as a function of the mean star rating of the choice set, for all choice sets (left panel) and broken down by choice set size (right panel). We observed a marked context effect in the Experiment, that participants were less likely to make a ratings-maximizing choice in sets with higher mean star ratings.

Analyzing choices, we observed participants were less significantly likely to choose the highest-rated ‘target’ option in choice sets with higher mean ratings (Figure 4B; *β*= −0.8538, 95% CI: −1.321, −0.4892, *p*<0.0001; Supplementary File 1G), but again, the variance of the star ratings of the choice set exerted no appreciable influence on ratings-maximizing choices (*β*= −0.8538, 95% CI: −1.321, −0.4892, *p*<0.0001). In other words, our laboratory experiment replicated the choice context effected observed in the real-world choice behavior—more valuable contexts were associated with less likely choices of the highest-rated option. Following our analysis of Yelp choices, we also probed for IIA violations in the 88% of choices made to the two best options in each choice set in the experiment, finding that—as we observed in the Yelp datasets— the average value of inferior distractor items significantly predicted participants’ tendency to choose the higher-valued of the two options (*β*=-0.6724, 95% CI: −0.8535, −0.5092, *p*<0.0001; Figure 4 – Figure supplement 1; Supplementary File 1H). Turning to expected satisfaction ratings, we observed a striking context effect on participants’ expected satisfaction ratings (Figure 5A), finding that more valuable contexts reduced participants’ expectations *(β=* −2.7376, 95% CI: −3.3713, −2.0962, *p*<0.001). This analysis, as in the real-world ratings, controlled for the influence of a number of variables that may have influenced participants’ choices, including the star rating and price of the chosen option (see Table 3).

**Figure 5.**
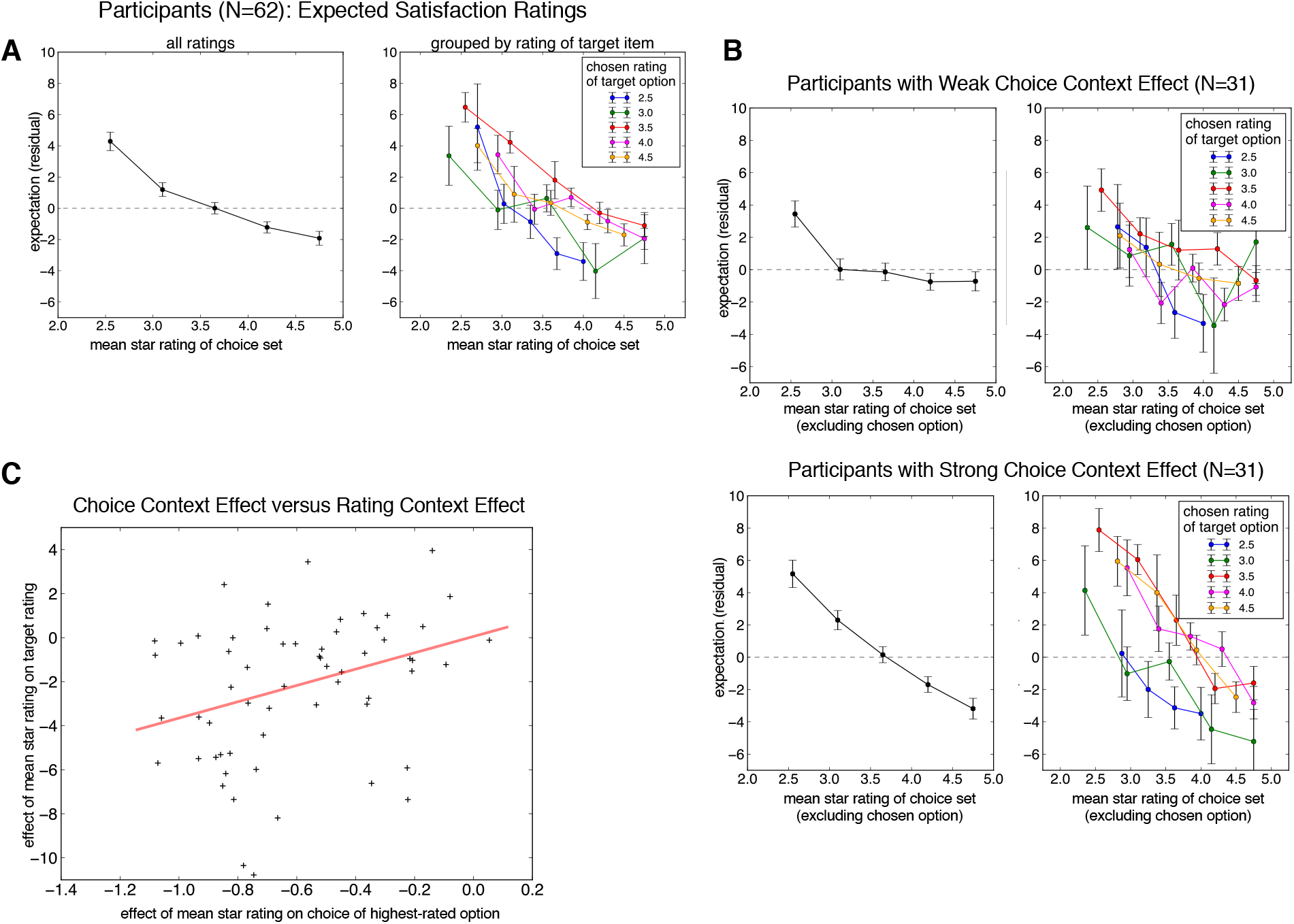
(A) Expected satisfaction ratings for the selected option, after controlling for a number of nuisance variables, as a function of the mean rating of the choice set, collapsed over all choice sets (left panel) and grouped by the rating of the selected option (right panel). We observed a marked context effect such that participants gave lower expected satisfaction ratings in higherrated choice sets. (B) Context effects for participants whose choice context effect sizes were weak (top row) and strong (bottom row). We observed that these rating normalization effects were stronger in participants with steeper normalization in choices. (C) Correlation between subjectlevel choice context effects (estimated from the group-level mixed effects regression) and the participant-level effect of mean rating of choice set (i.e., context effect with respect to ratings; Participants who exhibited stronger (negative) context effects in choice also exhibited a stronger (negative) context effect in expected satisfaction ratings, statistically confirmed by robust regression.

**Table 3.** Coefficient estimates for mixed-effects regression predicting expected satisfaction ratings as a function of the mean star rating of choice set excluding the selected option (i.e., the context effect), controlling for the rating of the selected option at the time of choice, log-transformed choice set size, the price of the selected option, and the log-transformed number of reviews of the selected option.

| *Coefficient* | *Estimate* | *95% CI* | *p-value* |
| --- | --- | --- | --- |
| (Intercept) | 29.3453 | 24.5856, 33.2665 | <0.0001* |
| mean star rating of chosen option | 12.078 | 11.5269, 12.5713 | <0.0001* |
| number of reviews for chosen option | 2.4053 | 2.0711, 2.6817 | <0.0001* |
| price of chosen option | 0.1306 | -0.4522, 0.703 | 0.336 |
| **mean star rating of choice set** | **-2.7376** | **-3.3713, -2.0962** | **<0.0001\*** |
| number of options (log) | -1.4998 | -1.9922, -0.9704 | <0.0001* |

In our account of the real-word ratings (Figure 3B), these (unobserved) contextually-computed expectancies are the basis of the comparison with a decision-maker’s true experience with the rated option. In our experiment, which examined choice and rating behavior in many of the same choice sets that Yelp users presumably faced in the real world, the observation that these expectations decrease sharply in accordance with the average value of the context supports our account of this comparison process: as contextually-bound expectations decrease, the user’s real-world rating—conceptualized as the true consumption utility minus expectations— increases. Put another way, these experimentally elicited expectations—which appear subject to contextual modulations of the surrounding choice set (Figure 5A)—provide a compelling explanation for the contextual effects observed in real-world rating data.

Finally, we probed the relationship between individual differences in the extent of context-dependence manifested in choice— quantified by the subject-level effect of mean choice set rating upon the probability of ratings-maximizing choice—and context-dependence evident in separately-measured expectation ratings. To do this, we examined context-dependence in expectation ratings separately for participants with small versus large context effects in choice (Figure 5B; via median split), finding that context dependence in ratings was markedly stronger in individuals whose choices also exhibited more context dependence. That is, the slope of the relationship between the mean value of the choice set (i.e., context) and the rated expectation was more pronounced for participants whose choices exhibited stronger contexts effects in choice—though, critically, these contexts effects on choice and ratings were observable at the aggregate, sample level. Further illustrating the concordance between context-dependence in choice and expectancy ratings, we found that the strength of context dependence in choice (quantified by the subject-level effect of mean rating of choice set upon ratings-maximizing choices) predicted the degree to which ratings were dependent on the average value of the choice set (Figure 5C; quantified by the subject-level random effect of mean rating of choice set; robust regression; *β*= 3.7073, *p*= 0.0104). In other words, the degree of context-dependence in an individual’s choices strongly related to the degree of context-dependence in their ratings, supporting the generality of these context effects to both choice and subjective valuation.

## Discussion

Here we examined whether real-world choices, evidenced in a large dataset of online restaurant reviews, are subject to context effects that have previously been observed in tightly-constrained laboratory settings (22, 23). We observed that ratings-maximizing choice behavior was systematically inflected by choice context, and importantly, this robust pattern of contextdependent choice replicated in two separate datasets which operationalize choice differently. We also found that subjective ratings were systematically biased by the rating distributions of the choice sets faced by decision-makers. This observed context-dependence appeared to be driven by the values (i.e., ratings) of the options alone, suggesting that this real-world rating information was evaluated—and acted upon, insofar as informing restaurant choices—in a relative and context-dependent, as opposed to absolute manner.

Further corroborating this account, in a laboratory experiment, we found evidence for context dependence in decision-makers’ choices and subjective expectations of options’ values, which, beyond replicating real-world choice context effects in a controlled setting, demonstrates that the value composition of a choice set can systematically alter decision-makers’ expectancy of an option’s value. Further, the concordance between the extent of context-dependence observed in individuals’ choices and context-dependence in ratings (Figure 6C) further supports the idea that contextually-computed subjective valuations—observable in expectancy ratings— serve as input to choices (23). Taken together, the marked context dependence we observe ‘in the wild’ and experimentally further buttress the idea that an option’s value is computed relative to its context, an idea pervasive across neuroscience, psychology, and behavioral economics (3, 5, 48). Computationally, relative—versus absolute—value representation holds the benefit that information processing resources are allocated efficiently to the encoding of subjective values (14, 15).

Our results raise a number of questions about the nature of these observed context effects. While our account postulates that the expectations informing the real-world ratings are computed in a relative, context-dependent fashion—which finds more direct support in our laboratory experiment (Figure 6A)—an open question concerns whether the subjective utility of consumption is also computed contextually, relative to the value composition of the choice set (or some other reference point). However, such effects could also arise from processes operating at consumption (or feedback, e.g. (49, 50)). Future work should examine, more directly, the extent to which subjective (consumption) utility might also exhibit context dependence.

While our analysis makes a core assumption that Yelp users’ choices (and ratings) are informed by ratings users observe on the Yelp website (or smartphone app) at the time of presumed choice, it may be the case that these online ratings may act here as a proxy for users’ knowledge about the value of options in the environment, rather than directly informing the observed choices. In other words, some users’ (absolute) valuations of options could be informed by direct experience with options or ‘word of mouth’ rather than star ratings. While user ratings are often a valid—if noisy—indicator of “true” quality of the options (51) (but see (40)), we remain agnostic about which source(s) of information are the basis for these choices. The qualitative concordance between real-world choice patterns (Figure 2A) and laboratory choices (5B), which were made in the absence of true knowledge about the options’ ‘true’ values—and instead, only on the basis of ratings information—hints that the rating information on Yelp might plausibly have been the basis for real-world users’ choices. Moreover, the low rates of ratings-maximize choice in the real-world choice data (in contrast to the experimental choice data) likely reflect other, possibly idiosyncratic considerations that decision-makers face in restaurant choice—for example, personal experience or other contextual factors not represented in the data, such as geographic, time, or social constraints.

It also is worth noting that our analysis approach—as in any real-world choice study (29, 31, 52)— relies critically on the operationalization of choice sets, which in our case were spatially-defined clusters of restaurants belonging to a single category. This simplified our analysis by guaranteeing that each restaurant appears in at most one choice set, and that restaurant options only competed with each other within the context of a single category. Of course, it is possible that an individual considers multiple restaurant categories when choosing between restaurants (31), or that, in choosing across different categories, a restaurant could present as an option in multiple choice sets (e.g., “pizza” and “Italian” restaurants). However, any such construction would presumably be orthogonal to the context effects observed here, absent any asymmetric relationship between restaurant rating and category membership. More directly, it is unlikely that these simplifying assumptions systematically hold the consequence that these robust context effects observed are the result of artifacts in light of the laboratorybased replication of these choice patterns. Nevertheless, future work could endeavor to understand how choice set rating distributions of different categories (and possibly even other choice sets of the same category) might bear on context-dependent choices and ratings. Finally, while a follow-up analysis revealed that these context effects were robust to the differing spatial definition of choice sets across the Yelp and Deliveroo datasets, we should note it is possible that individuals might, at times, strive to choose the a more globally “optimal” restaurant with respect to an entire city (or large region of a city) rather a single neighborhood per se.

Relatedly, it is possible that the set size effect we observe on value-maximizing choice— such that users were less likely to make an apparent ratings-maximizing choice in larger choice sets (Figure 2A) could be an artifact of our post-hoc construction of choice sets. That is, adding more options to the consideration set through any means will necessarily decrease the apparent likelihood of a decision-maker choosing any one option. Suggesting against this possibility, the same set size effects are observed in our laboratory data (Figure 5B) and have been previously observed by others [17,37]. Nonetheless, the context effects of interest here are the distribution of values (i.e., mean choice set rating) rather than the number of options in the set.

## Materials and Methods

### Analysis of Yelp Dataset

#### Data sources and selection of users and restaurants

Real-world data analysis focused on two data sources in the Yelp Open Dataset (obtained from https://www.yelp.com/dataset): 1) reviews of business, which consists of a unique user ID, a unique business ID, the date and time of review, the unique ID of the business reviewed, the user’s given star rating (ranging from 1 to 5 in whole numbers), and the text of the user’s review, and 2) data describing each business, which consists of unique business IDs, business names, GPS coordinates (latitude and longitude) and restaurant categories. We note that the reviews in this dataset reflects the output of Yelp’s recommendation software, which filters out suspect or biased reviews on the basis of quality, reliability, and user activity patterns.

We examined only restaurants in predominantly English-speaking US and Canada Metropolitan Statistical Areas (MSAs) with at least 1000 restaurants, leaving six US MSAs in the analysis: Charlotte-Concord-Gastonia NC-SC, Cleveland-Elyria OH, Las Vegas-Henderson-Paradise NV, Madison WI, Phoenix-Mesa-Scottsdale AZ, and Pittsburgh PA (US), and two Canadian MSAs: Calgary AB, and Toronto ON. We only examined choices pertaining to restaurant businesses—that is, all business entries containing the category “Restaurants” in the Yelp data, which totaled 59,315 restaurants in the US and Canada. Each business was mapped to an MSA with the US Census Bureau’s defined Core-Based Statistical Areas (in the US) or Census Metropolitan Areas (in Canada). We examined user reviews that were posted between January 1 2012, extending to November 14 2018 (the temporal end of the Yelp dataset).

#### Calculation of restaurant categories and neighborhood clusters

As business in the Yelp dataset are assigned multiple, unordered categories (e.g., “Restaurants, Italian, Pizza”), we inferred the single category that best described each restaurant. For each restaurant, we tabulated the occurrence of each Yelp-assigned category term in the text of the user reviews associated with that restaurant, and used the most frequently-occurring category term (e.g. “Pizza”). This ensured that each restaurant occurred in at most one possible consideration set, defined by restaurant category and neighborhood, preventing a restaurant from occurring in multiple consideration sets (e.g. both “Pizza” and “Italian”).

To define spatial neighborhoods within each MSA—which, alongside the restaurant categories, jointly defined the restaurant consideration sets—we employed the DBSCAN algorithm (35), a density-based spatial clustering algorithm that permitted us to define “clusters” based on users’ restaurant visit patterns, allowing neighborhoods of any arbitrary shape. The DBSCAN algorithm took the GPS coordinates of each restaurant review as input (e.g., 1,191,427 data points in the Phoenix-Mesa-Scottsdale, AZ MSA) and two parameters: a search radius (in decimal degrees), and a minimum number of data points within the search radius. Because the restaurant density and overall size of the MSAs considered vary considerably, the search radius and the minimum number of points were separately tuned for each MSA by perming nearest-neighbor analysis on the restaurants’ GPS coordinates and the total number of restaurants in each MSA, following previous work (36). Across all MSAs, the average number of restaurants, irrespective of category, per neighborhood (i.e., cluster) was 11.389 *(SD=* 1.346; see Supplementary File 1A for statistics describing resultant category and neighborhood sizes).

#### Selection of User Data

We only examined the choice and review behavior of users for whom there were at least 100 reviews in the Yelp dataset, yielding 4,591 unique users. Within each user’s series of reviews, we only considered restaurants in the most frequently-visited (i.e., rated) MSA for each user, for which we assume users likely have the most knowledge of the restaurant options. The resultant distribution of users per MSA was as follows: Phoenix-Mesa-Scottsdale AZ: 29.84%, Las Vegas-Henderson-Paradise NV: 28.33%, Toronto ON: 21.26%, Charlotte-Concord-Gastonia NC-SC: 7.382%, Pittsburgh PA: 4.849%, Cleveland-Elyria OH: 4.324%, Calgary AB: 2.078%, and Madison WI: 1.935%.

#### Calculation of Choice sets and Choice and Rating Measures

For every restaurant that a user reviewed, we analyzed that restaurant in the context of the restaurant’s category and its inferred geographic cluster —which, together, define a choice set. Our analysis only considered choices made within sets composed of 3 or more restaurants, with at least 3 unique ratings (the context effects of interest hold when we consider all possible choice sets), resulting in 53,118 choices examined in the final analysis. For each option in the choice consideration set, we computed the mean star rating for each option on the date of the user’s review—that is, considering reviews up to the point of that review but not including the review to capture the user’s possible informational state at the time of choice—rounding to the nearest half-star, as is seen by Yelp users (51). We computed the probability of choosing the (possibly non-unique) highest-rated restaurant in the choice set, following previous work (23, 24)—as a function of the mean star rating of the choice set in question computed over these rounded star ratings.

In our analyses of post-choice ratings, the Rating Deviation measure was computed as the difference between the mean rating of the chosen restaurant before that user’s rating and the user’s rating itself. To ensure user ratings measured pure context effects, the average star rating of the choice sets (the abscissa in Figure 4B) analyses omitted the chosen (i.e., rated) option. The residuals plotted in Figures 4B and Figures 6A-C are estimated from the mixed-effects regression models reported in Tables 2 and 3, omitting the mean star rating of the choice set as a predictor variable.

#### Replication Dataset: Yelp Check-ins

We analyzed the patterns of check-ins—time-stamped, geolocated record of a user having been at a particular business—as a proxy for visitations to restaurants in the Yelp dataset. Using the Yelp mobile app, a user can only “check into” a business, which will be shared with their Yelp social network (i.e. “friends”) if their current GPS coordinates are proximate to the coordinates of the business they are checking into. Unlike ratings, the aggregate number of check-ins to a business are not visible to other users, which mitigates the possibility that a checkin could be used to bias the rating distribution or popularity of a particular restaurant.

The Yelp dataset contains a list of timestamped check-ins for each business (check-ins per restaurant *M*=8,503; note these are not linked to specific users), which we analyzed similarly to the ratings-inferred choice analysis. For the 565,661 check-ins considered in the analysis, which spanned the years 2014-2016, we determined the surrounding choice set for that check-in using the same neighborhood and category definitions as in the ratings-based choice analysis and tabulated whether that check-in was made to the highest-rated option in that choice set. We then examined whether the mean star rating of the choice set influenced the probability of checking into the highest-rated option.

#### Replication Dataset: Deliveroo Orders

We analyzed choice patterns of customers ordering food on Deliveroo, an online food delivery service popular in several countries outside of North America (31). Our dataset consisted of 1,613,968 food orders, placed by 195,333 users ordering from 30,552 restaurants of 180 cuisine types across 197 cities, spanning February 2018 to March 2018. Similar to Yelp, each restaurant’s star rating (1-5 stars), the number of previous ratings for the restaurant at the time of the order, and the price of the restaurant. Here, we defined choice sets as unique combinations of cities and cuisine types (e.g., “Mexican” in Coventry, UK). We filtered out choice sets with less than 5 restaurants, leaving to 821 unique choice sets in total. We calculated the probability of a user ordering from the highest-rated restaurant, for each choice set separately over the course of the two months. Mirroring our analysis of the Yelp datasets, we then examined whether the mean star rating of each choice set influenced the user’s probability of ordering from the highest-rated restaurant in the set.

#### Inferential Statistics

In both the real-world and laboratory data, we estimated mixed-effects regression models, were using Markov chain Monte Carlo via the MCMCglmm package (version 2.32) for R (53), using logistic regression in models predicting choices of the highest-rated option with uninformative priors. Fixed effects, which are indicated in each regression table, were all taken as random effects over users (or participants). In all applicable models, we log-transformed the length of the text review, the number of reviews for the chosen option, and the number of options in the current choice set to mitigate skew apparent in these variables. In the experiment, subject-level effect sizes in choice and rating context effects (Figure 6) were computed as per-subject regression coefficients from the group analysis (conditioned on the group level estimates), superimposed on the estimated group-level effect.

### Choice Experiment

#### Participants

We collected data from 100 participants on Amazon Mechanical Turk (Crump et al., 2013), using the psiTurk package (54). This sample size, which we have adopted as a standard for all online studies employing within-subject designs, ensures adequate statistical power to detect meaningful differences across conditions, while at the same time protecting against false positives results or inflated effect sizes which can result from small samples. Further, this large sample size anticipates the participant exclusion rates, resulting from similar criteria, which we have observed in prior online investigations of this sort. Participants provided informed consent in accordance with the McGill University Research Ethics Board and were paid $4.00 USD for their participation. Thirty-eight participants were excluded from the sample due to failing to correctly respond to at least 60% of catch trials (see below), leaving a final sample size of 62 (age *M*= 41.88, *SD*= 12.00, 32% female). Note that the exclusion of these participants does not affect the significance of results reported above.

#### Design

Our experiment utilized 275 choice sets extracted from the real-world analysis described above. Each choice set was presented to participants in a 4×3 grid of restaurants depicting an image of each restaurant (from the yelp website), the average star rating of each restaurant (1 to 5 stars in increments of half stars, following the Yelp interface), the number of reviews used to compute the average star rating, the price of each restaurant represented as dollar signs ranging from one to four, and the category (i.e, cuisine the restaurant) (see Figure 5A). The size of each choice set varied from 3 (71% of the choice sets) to 7 (1%) options. Following our real-world analysis, restaurants in each choice were always drawn from the same category (e.g., ‘pizza’). Each participant viewed 275 choice sets, which were randomly divided into 135 choice trials, 135 rating trials, and 5 catch trials.

#### Procedure

Choice trials required participants to answer the prompt “Which restaurant would you choose if you were going to eat at a [category] restaurant?” among the available options. Participants indicated their choice by clicking on a cell in the grid with their mouse. On rating trials, a randomly chosen restaurant from the choice set was highlighted, and participants provided an expectation in response to the prompt “Imagine you ended up at [highlighted restaurant]. How satisfied do you think you would be with this restaurant if you were going to eat at a [cuisine type] restaurant?” Participants responded to this question using a slider bar that ranged from “Not satisfied” to “Very Satisfied” (the output of which was not visible to participants, but ranged from 0 to 100). Finally, each participant completed 5 catch trials (used to identify participants who did not follow instructions), in which they were required to choose the restaurant with the star highest rating. Importantly, no specific directions were provided concerning the information participants should use to make choices and ratings.

For all trial types, the available options were presented on the screen for 5 seconds during which time participants could not make a response, and following this viewing period, participants had 5 seconds to make a response. Following each choice and rating, participants rated their confidence in the previous response using a that ranged from 1 (“Not at all confident”) to 7 (“Very confident”). Participants had 5 seconds to make a confidence response. Confidence data were not examined in the present analyses.

## Acknowledgments

We thank Brendan Johns, Xi Liu, Jason Dolatshahi, and Nicholas Malecek for helpful discussions during the development of this work. This research was supported by an NSERC Discovery Grant, a CFI Infrastructure Grant to ARO, and a FRQNT Doctoral Fellowship to SD.

## Data Availability

Data from Yelp.com, anonymized experiment data, and analysis code can be accessed via the Open Science Framework at https://osf.io/ec5dx/ (DOI: 10.17605/OSF.IO/EC5DX). Due to access restrictions owing to the commercial and proprietary nature of the data, the Deliveroo data analyzed here was only accessible to one of the current study’s authors, and cannot be made publicly available. We are able to provide summary-level data (i.e. means depicted in Figure 2C) in tabular format upon request or upon publication. Researchers seeking access to this, or related raw data are encouraged to contact Deliveroo corporate.

## Supplementary Figure Legends

**Figure 2 – Figure Supplement 1.**
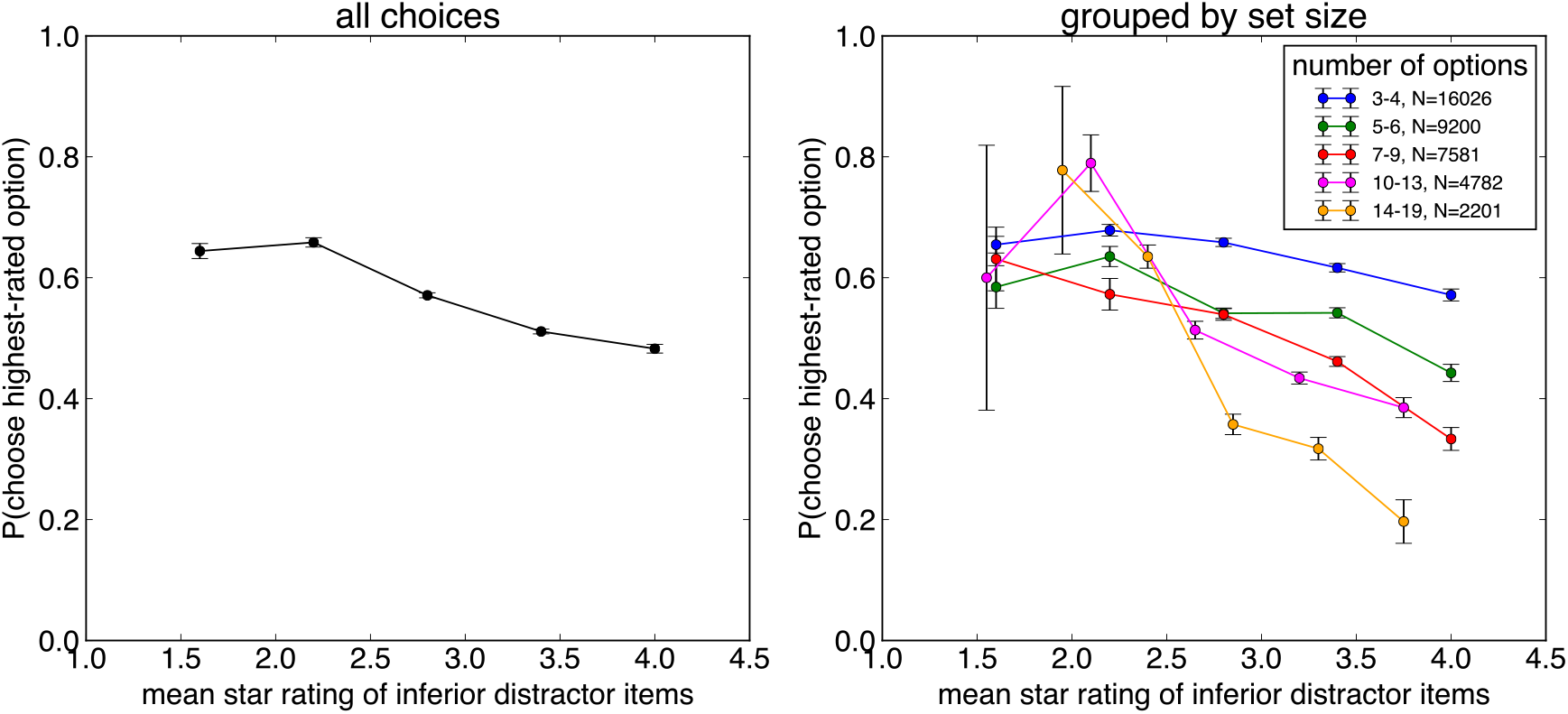
Proportion of choices to the highest-rated option, inferred from Yelp ratings conditioned on users choosing one of the two highest-rated options in the set, as a function of the mean star rating of the inferior “distractor” items (the sub-top-two-highest rated options), for all choice sets (left panel) and broken down by choice set size (right panel).

**Figure 2 – Figure Supplement 2.**
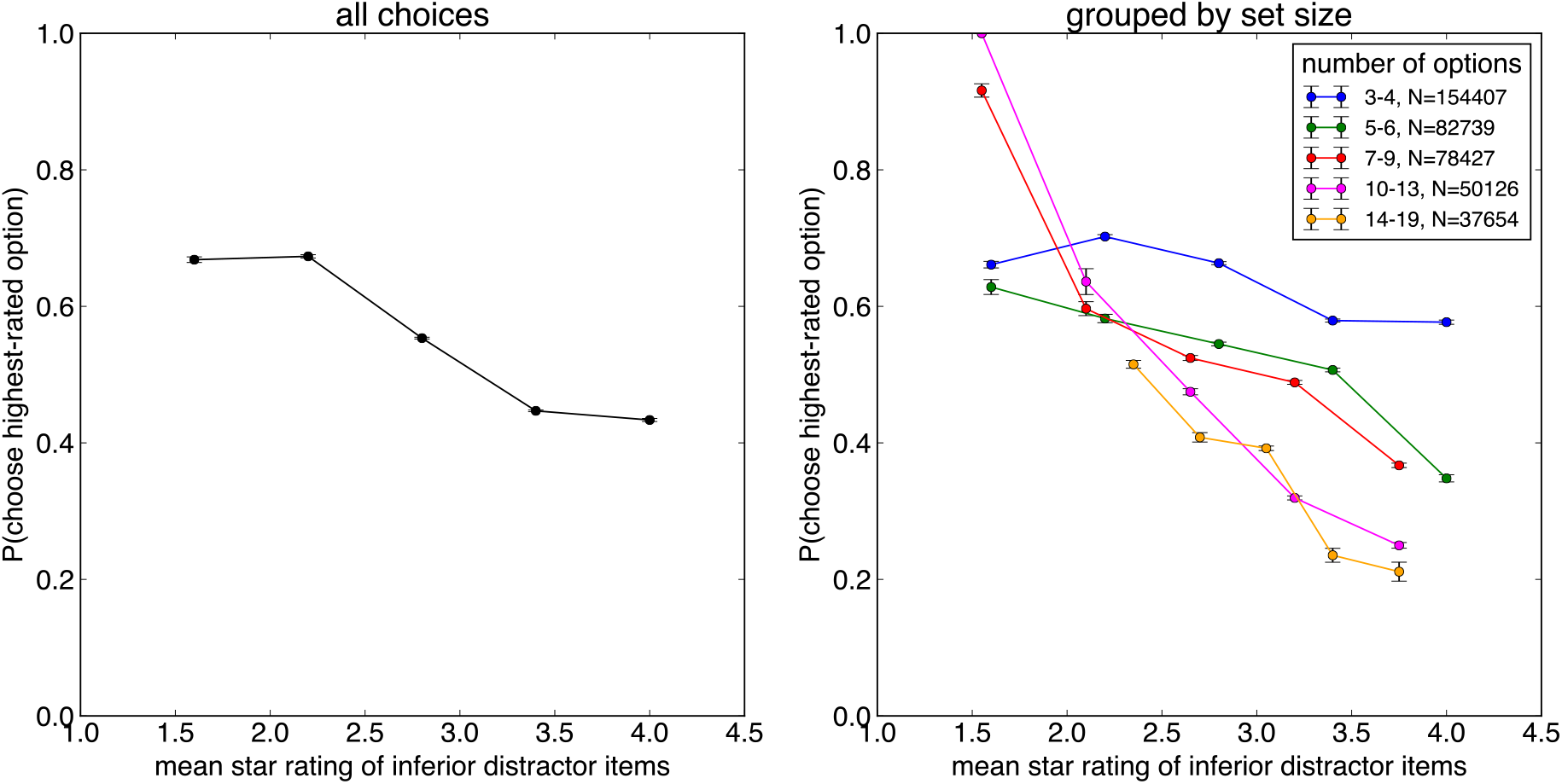
Proportion of choices to the highest-rated option, inferred from Yelp Check-ins conditioned on users choosing one of the two highest-rated options in the set, as a function of the mean star rating of the inferior “distractor” items (the sub-top-two-highest rated options), for all choice sets (left panel) and broken down by choice set size (right panel).

**Figure 4 – Figure Supplement 1.**
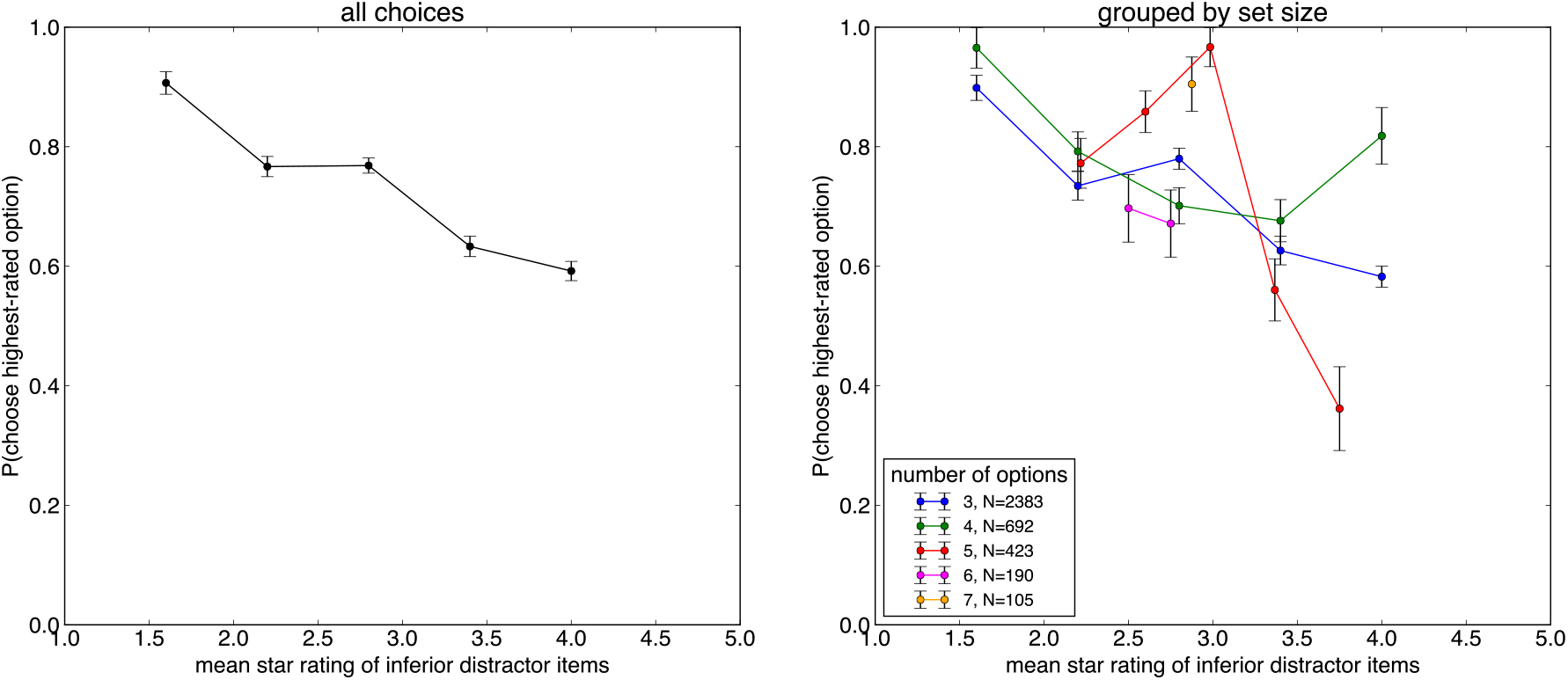
Proportion of choices to the highest-rated option in the Experiment, conditioned on users choosing one of the two highest-rated options in the set, as a function of the mean star rating of the inferior “distractor” items (the sub-top-two-highest rated options), for all choice sets (left panel) and broken down by choice set size (right panel).

